# Cyanophage-encoded lipid-desaturases: oceanic distribution, diversity and function

**DOI:** 10.1101/109157

**Authors:** Sheila Roitman, Ellen Hornung, José Flores-Uribe, Itai Sharon, Ivo Feussner, Oded Béjà

**Affiliations:** Faculty of Biology, Technion - Israel Institute of Technology, Haifa, Israel; University of Göttingen, Albrecht-von-Haller-Institute for Plant Sciences, Department of Plant Biochemistry, Göttingen, Germany; Migal Galilee Research Institute, Kiryat Shmona, 11016, Israel. Tel Hai College, Upper Galilee 12210, Israel; University of Göttingen, Göttingen Center for Molecular Biosciences (GZMB), Department of Plant Biochemistry, Göttingen, Germany.

## Abstract

Cyanobacteria are among the most abundant photosynthetic organisms in the oceans; viruses infecting cyanobacteria (cyanophages) can alter cyanobacterial populations, and therefore affect the local food web and global biochemical cycles. These phages carry auxiliary metabolic genes (AMGs), which rewire various metabolic pathways in the infected host cell, resulting in increased phage fitness. Coping with stress resulting from photodamage appears to be a central necessity of cyanophages, yet the overall mechanism is poorly understood. Here we report a novel, widespread cyanophage AMG, encoding a fatty acid desaturase (FAD), found in two genotypes with distinct geographical distribution. FADs are capable of modulating the fluidity of the host’s membrane, a fundamental stress response in living cells. We show that both viral fatty acid desaturases (vFADs) families are Δ9 lipid desaturases, catalyzing the desaturation at carbon 9 in C16 fatty acid chains. In addition, we present the first fatty acid profiling for marine cyanobacteria, which suggests a unique desaturation pathway of medium to long chain fatty acids no longer than C16, in accordance to the vFADs activity. Our findings suggest that cyanophages fiddle with the infected host’s cell, leading to increased photoprotection and potentially enhancing viral-encoded photosynthetic proteins, resulting in a new viral metabolic network.

## Introduction

Viruses are the most abundant entity in the oceans, yet the vast majority remains uncultured^1-4^. Cells lysed by viruses contribute to energy and nutrients flux in the oceans, while infected cells could also affect global biogeochemical cycles^5-9^. Viruses carry in their genomes a wide variety of Auxiliary Metabolic Genes (AMGs), capable of complementing or redirecting the infected host metabolism resulting in increased viral fitness^10^. Cyanophages, phages infecting marine cyanobacteria, display a broad array of AMGs, including photosynthetic light reactions components^11-20^. Photosystem-I (PSI) genes in cyanophages (vPSI) are found in two main genotypes, arranged in cassettes of seven (*psaJF,C,A,B,K,E,D*) and four (*psaD,C,A,B*) genes, dubbed vPSI-7 and vPSI-4, respectively^13,21,22^. Since there are no cultured representatives of vPSI-4 phages, little is known regarding their potential influence on the infected host metabolic capacities.

Several AMGs are potentially involved in photoprotection of the infected cyanobacterial cell. For example, high light inducible proteins enable the dissipation of excess light energy and the correct functioning of the photosynthetic light reactions^23^, and are widely found in cyanophages^12,14,24^. Photosystem II (PSII) reaction center protein D1 (encoded by the *psbA* gene), was shown to be constantly damaged during photosynthetic activity and must be repaired and *de-novo* synthesized in order to maintain photosynthesis active^25^. The viral *psbA* gene is expressed upon infection^26-28^ and it was suggested to increase phage fitness^29,30^. In addition, many cyanophages carry genes for a plastoquinol terminal reductase (PTOX), potentially involved in photoprotection of PSII^19,20,31^. Based in the accumulating data in cyanophages AMGs *repertoire*, it appears that photoprotection of the cell is a central need of the infected cell, the “virocell”^32^ metabolism.

Another, rather unexplored mechanism for coping with photoinhibition in cyanobacteria includes the desaturation of the membranes lipids; polyunsaturated fatty acids are critical for growth and for photoinhibition tolerance of cyanobacterial cells^33-37^. Membrane fluidity also affects the activation of D1 and its *de novo* synthesis, leading to a higher recovery rate of PSII activity^33^. In cyanobacteria, lipids desaturation is performed at fatty acid residues esterified to a glycerolipid by membrane-bound acyl-lipid front-end desaturases (*des*), associated with cytoplasmic and thylakoid membranes. Molecular oxygen and an electron donor (ferredoxin) are required for fatty acid desaturases (FADs) activity^37-39^. Four *des* genes can be found in cyanobacteria, encoding for DesA, DesB, DesC and DesD, catalyzing the desaturation at carbon Δ12, Δ15, Δ9 and Δ6 (counting from the carboxy group), respectively^34,40-42^. Cyanobacteria have been classified into four groups based on their fatty acid desaturation degree, depending on the length of their fatty acids (C16 or C18), the amount of the double bonds (zero to four per fatty acid chain), and the *sn* position of the desaturated fatty acid (*sn-1* and/or *sn-2* at the glycerol backbone)^43^. However, marine unicellular cyanobacteria, namely *Synechococcus* and *Prochlorococcus* do not fit in any of the four classic groups based on their FADs composition, carrying only *desC* and *desA* genes^44^. DesC performs the first desaturation of fatty acids at position Δ9 and is present in all cyanobacterial strains^43,44^. These monounsaturated fatty acids are essential for growth. Consequently, *desC* knockout mutants must be supplemented with unsaturated fatty acids in order to survive^36,45^.

Here, we report the identification and characterization of two novel and widespread cyanophage-encoded FADs (vFADs) families. The vFADs were expressed using a heterologous yeast system and were identified as DesC-like FADs, catalyzing the desaturation at carbon Δ9 in C16 fatty acid chains. In addition, we performed, to our knowledge, the first fatty acid analysis of marine cyanobacteria and found their lipid composition to be unique among cyanobacteria. Our results suggest that marine cyanobacteria have a rare pathway for fatty acid desaturation, and phages desaturases are well suited to fit in.

## Results and discussion

To enrich our knowledge regarding uncultured cyanophages carrying photosynthetic genes we conducted a metagenomic survey in a re-assembly (Sharon et al., *in prep*) of the microbiome^46^ and virome^1^ datasets from the *Tara* Oceans expedition, a comprehensive sampling project of oceanic microbial diversity. Using the sequence of a viral PSI psaD gene as query for TBLASTX, we identified a 64 kbp contig containing a vPSI-4 cassette in the assembly of station 70 (South Atlantic Ocean, File S1). The contig was extended up to 94 kbp with recruitment of reads from the same station. This contig is predicted to have originated from a cyanophage of the *Myoviridae* family (T4-like phages), based on RegA (Fig. 1a) and Transaldolases (Fig. 1b) maximum likelihood phylogenetic protein trees, and the presence of three tRNA genes (Fig. 2) widely found among cyanomyophages^47^. The contig contains structural and DNA replication genes, along with various AMGs common in cyanophages, such as *talC*^24,48^, peptide deformylase^18^, *psbA* and *psbD*^11,26^, ferredoxin^24,48^, *phoH*^49^, among others (Fig. 2, Fig. S1, Table S1). Surprisingly, we also identified a gene coding for a putative vFAD, this being the first report of a cyanophage potentially interfering with fatty acid metabolism in the infected host cell. Using the identified vFAD gene sequence we were able to retrieve 139 contigs containing vFADs among various viral genes (File S1) from publicly available metagenomic datasets (Table S2).

**Figure 1:**
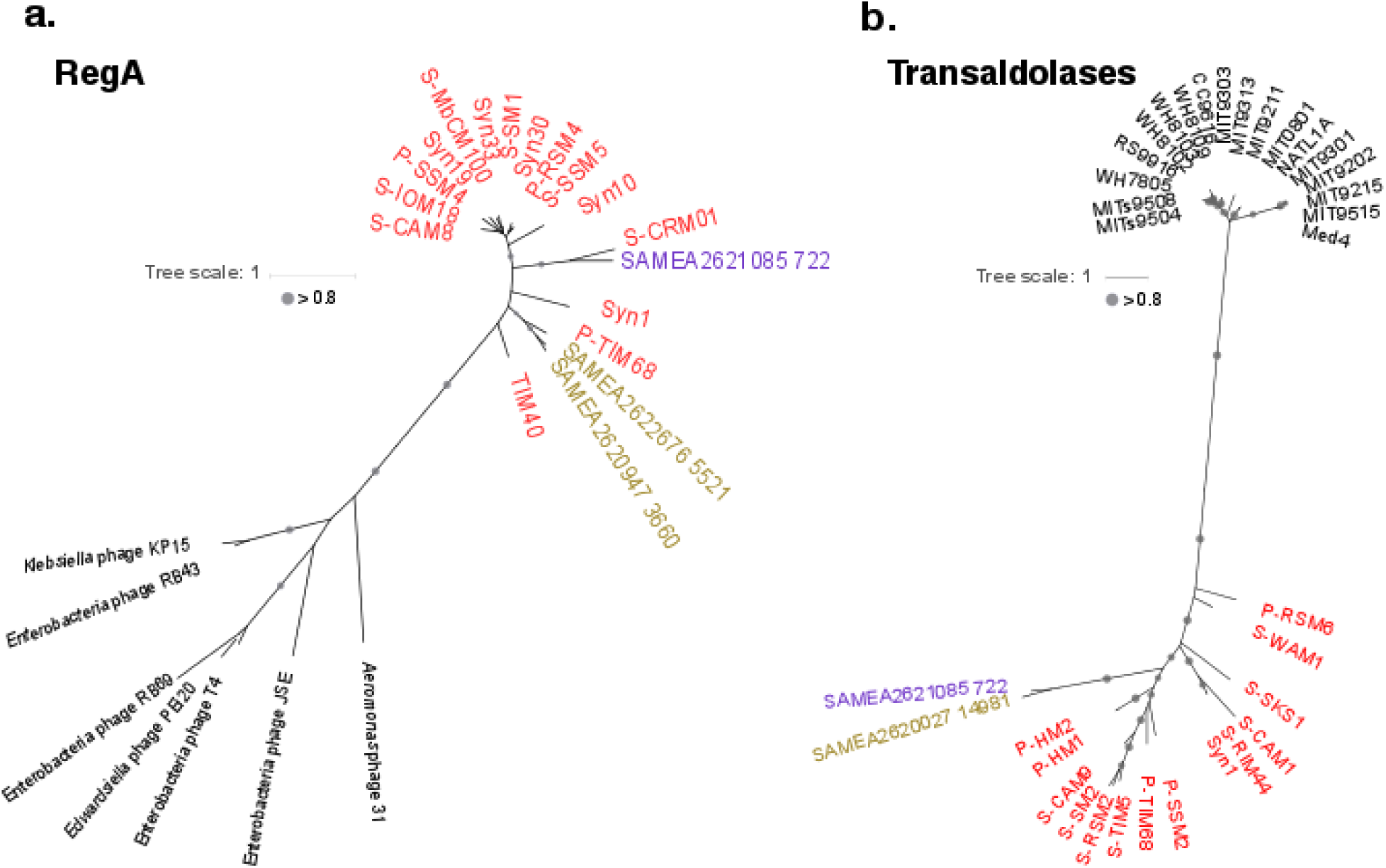
**Maximum likelihood phylogenetic trees of a.** RegA and **b.** Transaldolases (Tal A/B and TalC). Circles represent bootstrap values higher than 0.8. Cyanophages are colored in red, other bacteriophages are depicted in black. The RegA and TalC found in the 94 kbp contig are marked in purple. Contig names in gold originate in contigs retrieved in this project carrying both DesC and RegA/ DesC and TalC, clustering within the vFAD-I family. The scale bar indicates the average number of amino-acid substitutions per site.

**Figure 2.**
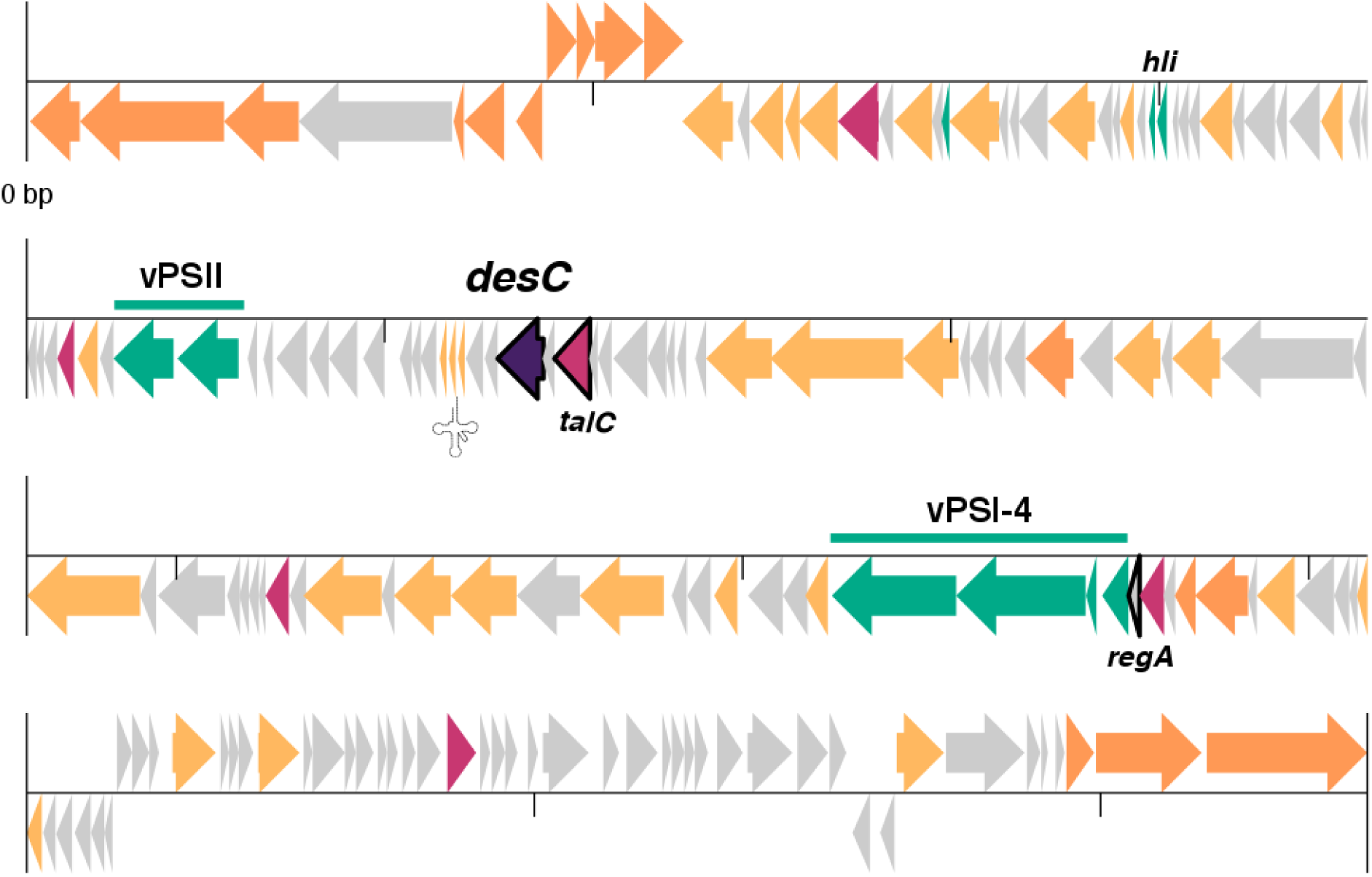
**94 kbp contig genomic map.** Grey arrows represent hypothetical and conserved hypothetical proteins. Orange arrows are virion structural and packaging genes. Yellow arrows stand for genes encoding DNA replication and metabolism modification proteins. Auxiliary Metabolic Genes are depicted in pink, while AMGs related to photosynthesis are colored in green. The fatty acid desaturase gene is colored purple. Genes encoding proteins used in phylogenetic trees in Fig. 1 (*regA, talC*) and Fig. 3 (*desC*) are contoured in black. Three tRNA genes are marked with a single tRNA icon. A detailed figure and list of open reading frames can be found in Fig. S1 and Table S1, respectively. Open reading frames sequences can be found in File S2.

The vFAD gene encodes for a putative acyl-lipid desaturase, a membrane-bound enzyme that catalyzes the front-end desaturation of fatty acids esterified to glycerolipids. The protein is homologous to membrane-bound DesC Δ9–front-end desaturases found in cyanobacteria (and plants) (Fig. 3). Moreover, it contains the three characteristic Histidine motifs of DesC-like desaturases, two HXXXHH and a HXXXXH, potential ligands of di-iron center in the active site of the enzyme^43^. Interestingly, some *Synechococcus* and *Prochlorococcus* strains (mainly low light adapted, clade IV) have two types of DesC (*Synechococcus* strain CB0101 has three) and these proteins cluster separately into two different branches in the DesC phylogenetic tree (Fig. 3), indicating a possible specialization for each type. Since the marine cyanobacterial FADs (cFADs) specific activity is yet unknown, we will refer to them as cFADs Δ9-3 and Δ9-4, according to the classification given by Chi *et al*.^44^. However, some *Prochlorococcus* strains carry a single DesC corresponding to Δ9-4 (cFAD Δ9-4 group in Fig. 3), while some *Synechococcus* strains contain only one Δ9-3 protein (cFAD Δ9-3 in Fig. 3). Accordingly, vFADs can be found in two genotypes, forming monophyletic branches in the phylogenetic tree. These groups correspond to the unicellular marine cyanobacterial types, although they share less than 70% identity on the protein level, thus we decided to denominate them vFAD–I (Fig. 3, shaded gold) and vFAD–II (Fig. 3, shaded purple). We retrieved more vFADs-I contigs than vFADs-II from the metagenomic datasets analyzed, however the first vFAD discovered, found in the 94 kbp contig, clusters within family II (marked with a black arrow in Fig. 3).

**Figure 3:**
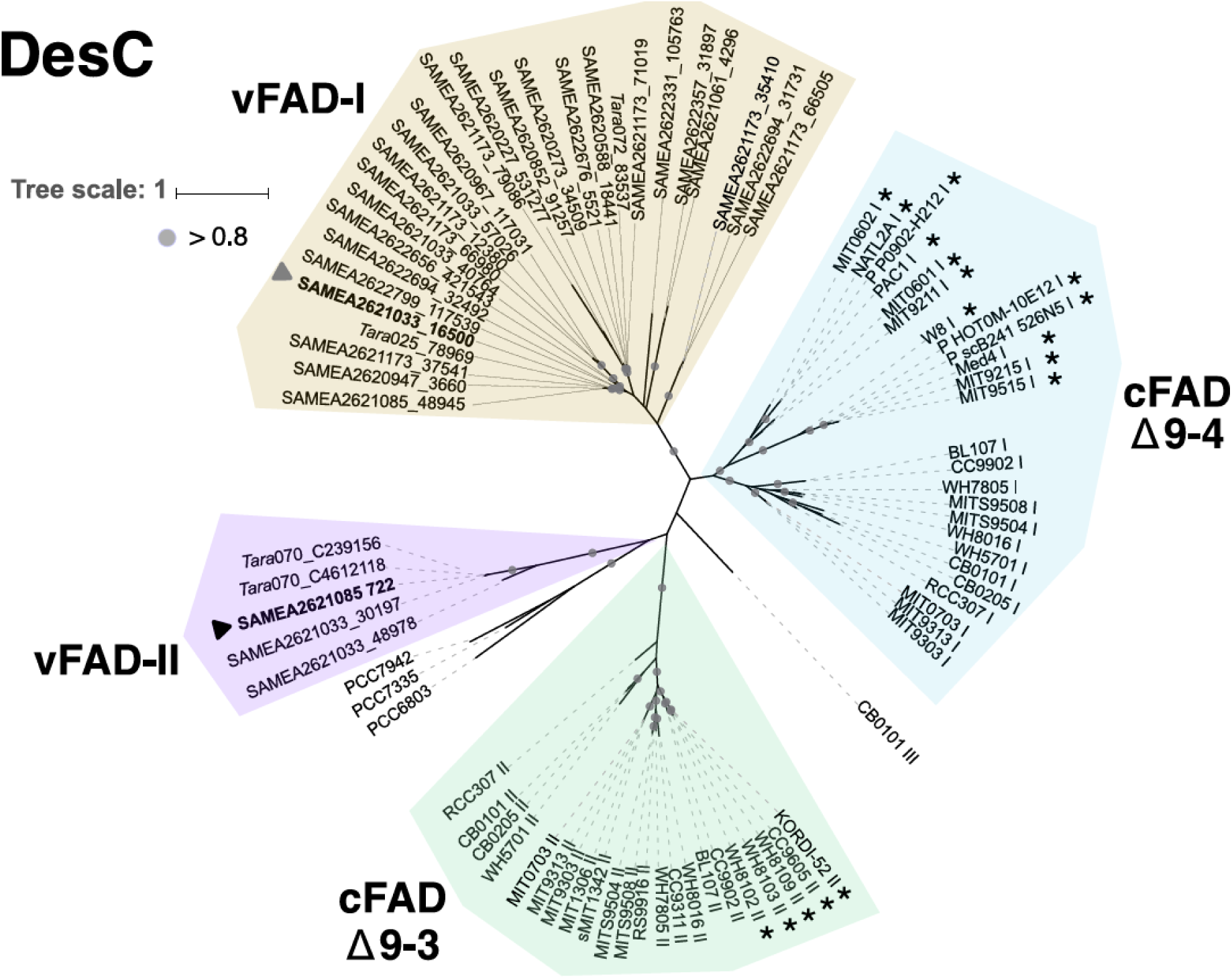
**Maximum likelihood phylogenetic tree of DesC.** Circles represent bootstrap values higher than 0.8. Viral fatty acid desaturases classified as families I and II are shaded in gold and purple, respectively. Cyanobacterial desaturases (cFADs) are shaded green and blue for Δ9-3 and Δ9-4 groups respectively. Black and grey arrows indicate the sequences chosen for expression in yeast. Stars indicate picocyanobacterial strains carrying only one *desC* gene. The scale bar indicates the average number of amino-acid substitutions per site.

vFADs families show distinct biogeography (Fig. 4a). vFADs-I are widespread in the oceans (Fig. 4a, golden dots), being found all along the Pacific and Atlantic Oceans, the Indian Ocean and the Mediterranean Sea. In contrast, vFADs-II are present only in the Southern Pacific and Southern Atlantic Oceans as well as in the Indian Ocean (Fig. 4a, purple dots). Interestingly, the geographical distribution and abundance of vFADs-II is similar to the data found for uncultured phages carrying the vPSI-4 gene cassette^22^, which is also found in the 94 kbp contig (Fig. 2). To assess the vFADs relative abundance we mapped the raw reads from the *Tara* Oceans microbiome and virome to the vFADs identified in the metagenomics survey. In sampling stations where both vFADs families were present, vFADs-I were more abundant by at least one order of magnitude than vFADs-II (Fig. 4b, Fig. S2). It is worth noticing that even when most of the reads mapped to the vFADs originated from the virome datasets, probably originating from free viral particles (Fig. S2), we also found many reads in the microbiome datasets, in agreement with the findings of Philosof *et al.*, that reads of cyanophage origin are commonly found in the microbial fractions of marine samples^50^. In comparison, reads mapping to cFADs originated almost exclusively from the microbiome datasets (Fig. S2).

**Figure 4.**
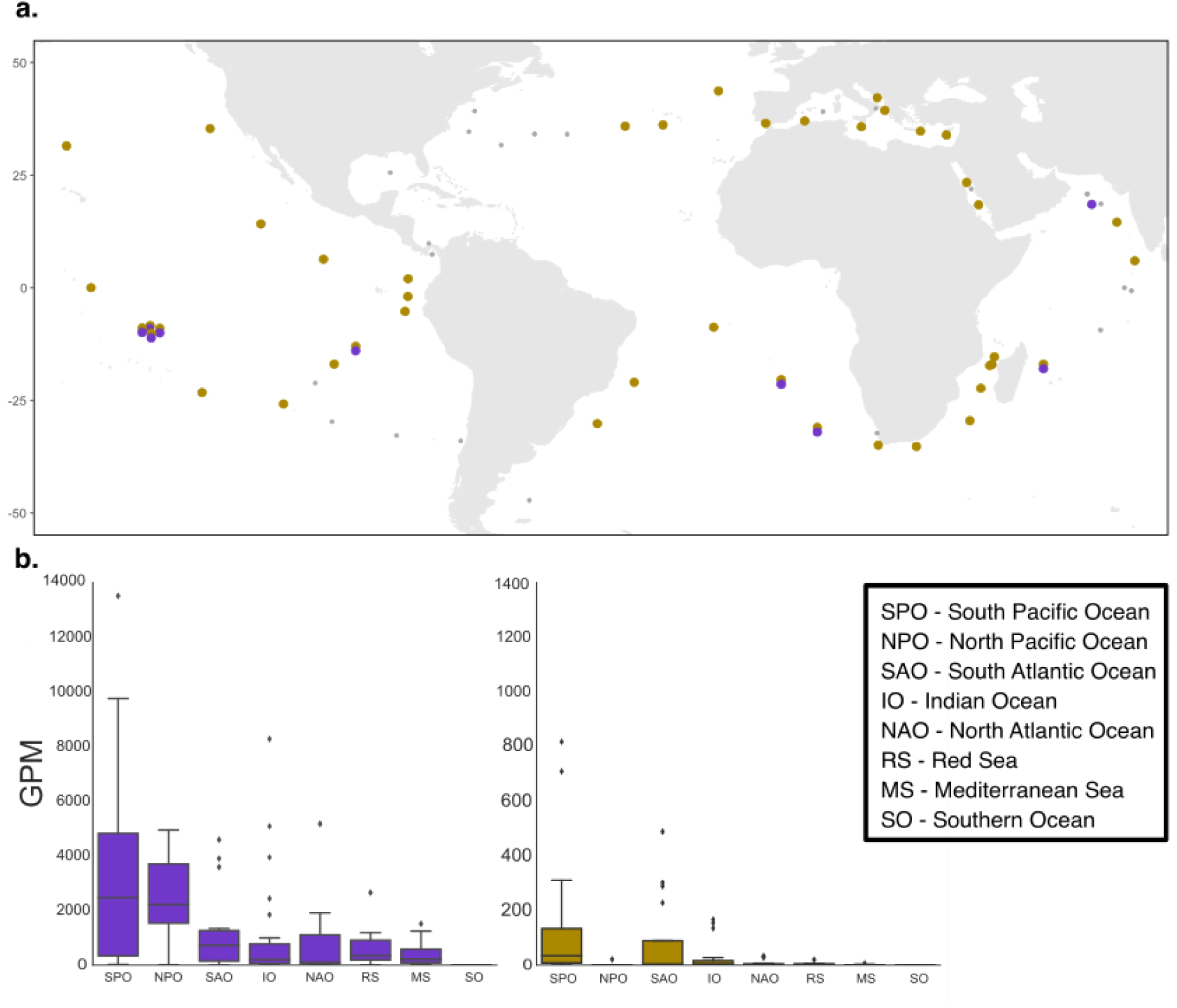
**a. Map of *Tara* Oceans stations analyzed in this project.** Gold dots represent stations positive for vFAD-I reads; purple dots mark stations positive for vFAD-II reads. Grey dots stand for stations were no reads for vFADs were found. Latitudes are marked at the left of the map. **b.** Relative abundance of vFADs from families I and II (depicted in purple and gold respectively). GPM – Genes per million.

In order to confirm the vFADs activity we expressed the viral genes in a heterologous system using the *Saccharomyces cerevisiae* strains INVSc1 and the FAD mutant Ole1^51^. While the INVSc1 strain contains mono-unsaturated (at position Δ9) and saturated long chain C16 and C18 fatty acids, the Ole1 mutant strain features only saturated fatty acids and has to be supplemented with unsaturated fatty acids for normal growth. The lipid profile of INVSc1 cells expressing vFADs could not be distinguished from cells transformed with an empty vector, suggesting for a possible Δ9 desaturation activity. This was confirmed by lipid profiles of Ole 1 mutant strains expressing vFADs (Fig. 5). Both vFADs show Δ9 desaturase activity, acting specifically on C16 chains of lipids in yeast. Marine cyanobacteria show a unique pathway for acyl-lipid desaturation, among cyanobacteria, containing only *desC* and *desA* genes for desaturation of carbons Δ9 and Δ12, respectively^44^, that may start with the C16 fatty acid (palmitic acid); yet their lipid profiles were not yet determined. We therefore performed fatty acid profiling of three cyanobacterial strains: *Prochlorococcus* Med4, carrying a cFAD Δ9-4, *Synechococcus* WH8109, carrying a cFAD Δ9-3, and *Synechococcus* WH7805 carrying both (Fig. 6a,b). All three strains contain a specific fatty acid profile, that interestingly, shows an unusual large amount of C14, compared to profiles of freshwater cyanobacteria^52^, and a very low amount of C18 fatty acids.

**Figure 5.**
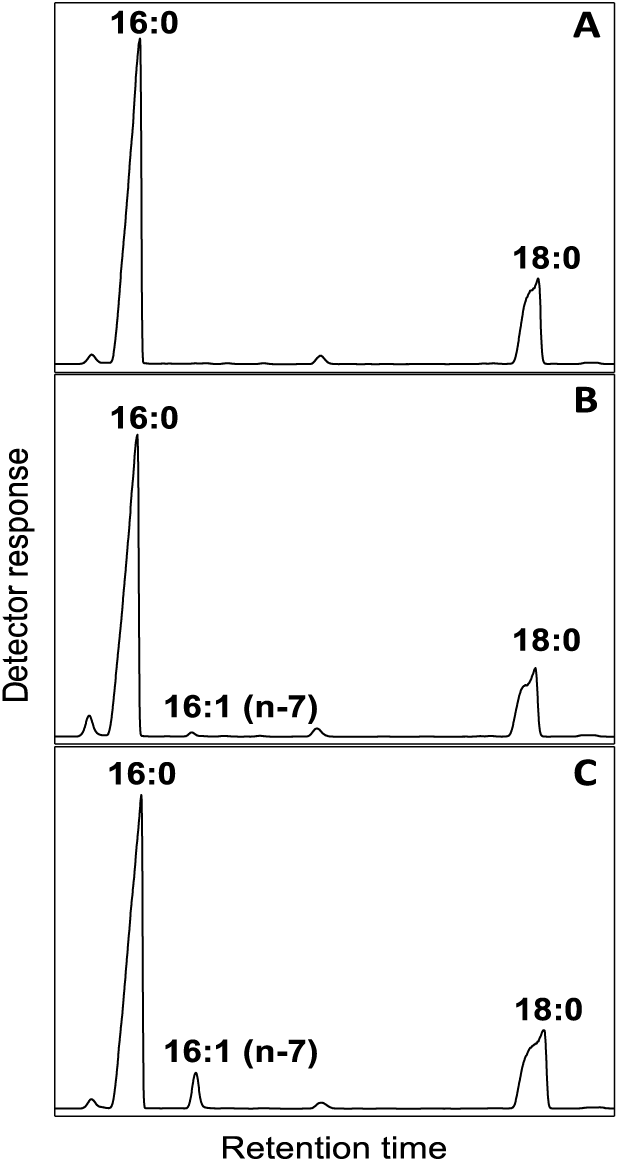
**GC/FID analysis of FAMEs isolated from Ole1 yeast cells expressing vFADs.** After lyophilization the esterified fatty acids were transesterified with sodium methoxide and analyzed by GC/FID (see methods). **A**. Chromatogram of the control yeast, transformed with an empty pYES2/CT vector. **B**. Chromatogram of the Ole1 yeast expressing vFAD-I (marked with a grey arrow in Fig. 3). **C**. Chromatogram of the Ole1 yeast expressing vFAD-II (marked with a black arrow in Fig. 3).

**Figure 6.**
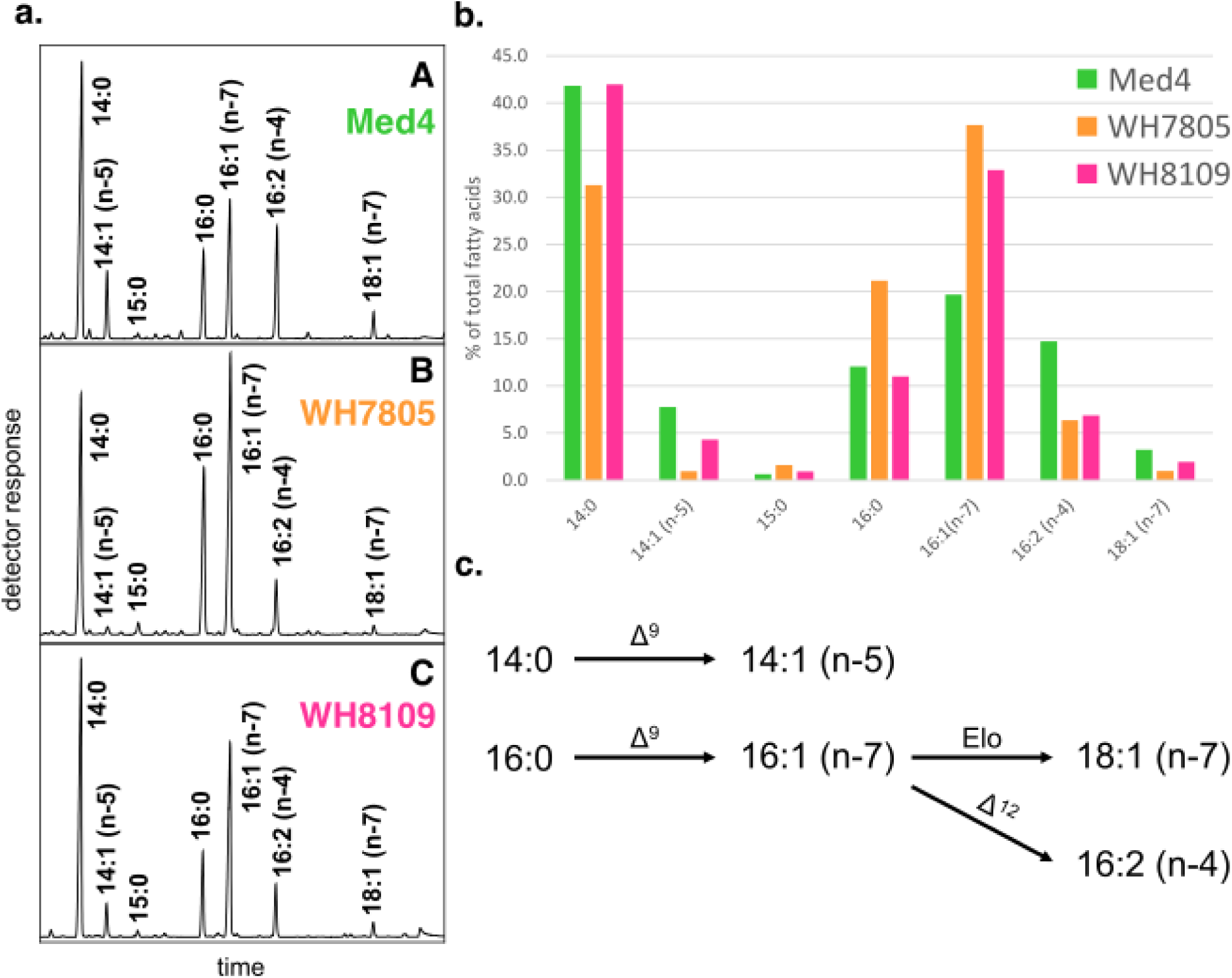
**a. GC/FID analysis of FAMEs isolated from cyanobacteria.** FAME were prepared from lyophilized cells using acidic methanolysis, and analyzed by GC/FID (see methods). Position of double bonds was verified by GC/MS analysis, after converting FAME to DMOX derivatives (see File S3). A – *Prochlorococcus* Med4, B- *Synechococcus* WH7805, C- *Synechococcus* WH8109. **b.** Fatty acids profile of the marine cyanobacterial strains. Fatty acids are expressed as the percentage of total fatty acids. Green bars – *Prochlorococcus* Med4; orange bars – *Synechococcus* WH7805; pink bars – *Synechococcus* WH8109. **c.** Pathway scheme for the biosynthesis of fatty acids in the analyzed cyanobacteria. *De novo* synthesis ends either with carbon chain length 14 or 16 yielding 14:0 and 16:0, respectively. Next, these fatty acids may be desaturated by a DesC-type Δ9 desaturases yielding 14:1 (n-5) and 16:1 (n-7), respectively. The later may then be further elongated (Elo) into 18:1 (n-7) or again be desaturated by DesA-type Δ12 desaturase yielding 16:2 (n-4).

Both, the heterologous expression as well as the fatty acid profiles of the marine cyanobacteria, hint to an unusual substrate specificity of the picocyanobacterial and viral desaturases (Fig. 6c). The recombinant vFAD enzymes only used C16 fatty acid chains as substrate but not C18 (C14 is almost not available in the used yeast strain), while cyanobacteria fatty acid profiles display desaturation at the Δ9 carbon for C14 (n-5) and C16 (n-7) but not in C18 fatty acid chains (Fig. 6a). Monounsaturated C18 (n-7) contains the double bond at position Δ11, thus being the result from elongation of monounsaturated C16 and not of *de novo* desaturation of saturated C18. We propose that viral and picocyanobacterial desaturases have special substrate specificity towards fatty acid chains no longer than C16. However, we cannot discard the possibility that there is little or no synthesis of C18 fatty acid chains in these cyanobacterial strains, thus the lack of substrate could explain their unusual specificity.

Based on the vFADs activity assay and the fatty acid profile of marine cyanobacteria, we devised a model for cyanophage fatty acid desaturases activity (Fig. 7). Viruses infecting eukaryotic organisms carry fatty acid metabolism AMGs for the synthesis of their unique outer lipid envelope^53,54^, lysing the host’s cell^54^ or to enable the replication of their genome^55^. Cyanophages, on the other hand, might carry vFADs in order to cope with stress resulting from photoinhibition and maintain homeostasis of the virocell during infection. DesC is essential for life; from all four desaturases found in cyanobacteria, only DesC is constitutively expressed^56,57^, maintaining its activity regardless of the light/dark cycle^37,56^. Moreover, it has been shown that the introduction of the first double bond in membrane lipids (Δ9 desaturation) has the most significant effect on the fluidity of the membrane, while subsequent desaturation steps have a smaller impact^57,58^. Desaturation of 16:0 fatty acids to 16:1 was also shown to occur within 10 hrs after cold stress induction without *de novo* synthesis of fatty acids^37^, meaning that changes in the cell’s environment can trigger an immediate stress response by modulating the membrane’s fluidity. However, *de novo* synthesis of desaturases is necessary^37^, as *desC* mRNA half-decay time was shown to be of approximately 10 minutes^57^, and a marked decrease in *desC* mRNA copies was shown after three hours in viral infected *Synechococcus sp.* WH7803 cells^59^. Since cyanophages infection can last several hours^26,27,60^, it might be beneficial for the virocell to maintain or even increase the fluidity of the membranes to cope with stress and photoinhibition. In the 94 kbp contig, we found along with the vFAD, vPSII and vPSI genes, whose activity might benefit from modifications in membrane fluidity, and a gene encoding for ferredoxin, that could potentially act as the electron donor to the vFAD.

**Figure 7.**
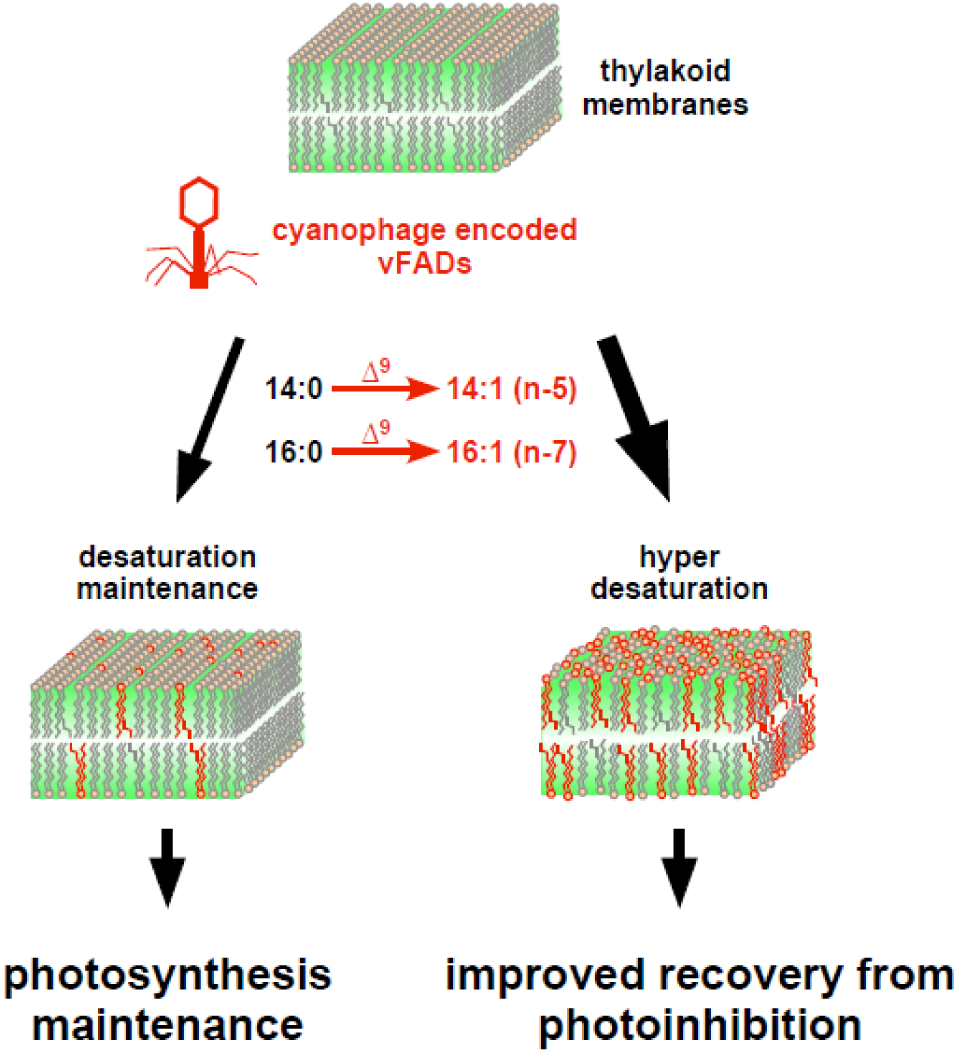
**Model for vFADs activity.** Upon infection, phages carrying vFAD genes can increase the desaturation level of pre-existing saturated C14 and C16 fatty acid chains, leading to higher membrane fluidity that might lead to improved recovery from photoinhibition. Maintaining and increasing the fluidity of the thylakoid membranes could affect the photosynthetic performance of the infected host cell. Both outcomes may lead to a higher fitness of the phage.

Marine *Synechococcus* and *Prochlorococcus* are among the most abundant photosynthetic organisms on Earth, and it was estimated that cyanophages lyse between 0.005 to 10% of cyanobacteria daily^61,62^. During infection, the virocell’s physiology is remarkably different from the original, uninfected cyanobacteria, as phages bring new metabolic capabilities with potential to rewire the host’s metabolism. Here we report a novel pathway in cyanophages, *i.e.* fatty acid metabolism that could have an overall impact on the virocell’s performance. This might lead to a higher fitness of the phage and to a change in the quality of the debris left after burst, which becomes part of the dissolved organic matter (DOM) used by heterotrophs and it is shunted back into the food web^7^. As we keep unveiling rare phage capabilities, we realize that their roles in the environment are far greater than expected.

## Methods

***Metagenomic data analysis.*** Metagenomic datasets from the *Tara* Oceans microbiome^46^ and virome^1^ were re-assembled (Sharon et al., in prep) using IDBA-UD^63^ assembler providing higher quantity of longer scaffolds than previously reported^46^. Errors in the assembly were corrected using two read-mapping based in-house tools (Sharon et al., in prep). Viral *psaD* sequences obtained in a previous study of vPSI-4 genes^22^ were used as query to recruit scaffolds in the re-assembled *Tara* Ocean dataset using TBLASTX^64,65^ with the default parameters. One of the identified scaffolds, SAMEA2621085 (station 70, depth 5m, 0-0.22 filter), contains the four genes of vPSI-4 (*psaD,C,A,B*). The scaffold carrying the vPSI-4 genes was extended using the mini-assembly technique described in^66^. This process lead to the recruitment of other fragments of the same genome until no further elongation could be reached. The resulting 94 kbp fragment went through QC, and consistency of the extended scaffold was confirmed by mapping the sample reads to the scaffold using Bowtie2^67^.

ORFs were identified in the 94 kbp contig using GeneMark^68,69^ and manually annotated using BLASTX (default parameters). The vFAD protein sequence was used as query for a TBLASTN search (e-value 0.1) against metagenomic datasets (Table S2). All retrieved contigs were screened using BLASTX (e-value 10e-10) against the NCBI non-redundant (nr) protein database to identify all putative proteins in the contigs. Fatty acid desaturases from cyanophage origin were selected based on top hits with less than 70% identity to picocyanobacteria.

Relative abundance of vFADs was calculated using kallisto^70^. A collection of DNA sequences (Table S3) composed of bacterial cFADs and the BLASTX identified vFADs was used to build a kallisto-index using default parameters. The kallisto-index was used for the pseudo-alignment of the *Tara* Oceans microbiome and virome reads. The “estimated counts” (est_counts) and “effective length”(eff_lenght) obtained from the kallisto output for all the analyzed datasets were used to calculate gene relative abundance in genes per million (GPM) in a similar manner to the calculation of transcripts per million (TPM)^71^.

***Phylogenetic construction and analysis.*** Newly identified vFADs, *talC*, and *regA* gene sequences were translated to proteins according to the correct open reading frame and aligned along with sequences from picocyanobacteria and cyanophages retrieved from GenBank. Multiple sequence alignments were created using ClustalX v2.1^72^. Maximum likelihood phylogenetic trees were constructed using the phylogeny.fr pipeline^73^, including the PhyML v3.0^74^ and the WAG substitution model for amino acids^75^. One hundred bootstrap replicates were performed for each analysis. See Files S4-S6 for the alignments used to construct the trees.

***Expression of vFADs.*** One representative from each of the vFADs families (SAMEA2621033_16500 for vFADs-I and SAMEA2621085_722 for vFADs-II, marked with a grey and a black arrow, respectively in Fig. 3) were chosen for expression. We performed codon usage (CU) adaptation for optimal expression in yeast using Integrated DNA Technologies (IDT) tool for codon optimization to *Saccharomyces cerevisiae* CU. DNA fragments, as gBlocks Gene Fragments (IDT), were cloned into the pYES2/CT vector (Thermo Fisher Scientific) and sequenced to confirm their identity. The plasmids were transformed into yeast strains INVSc1 and Ole1 (*ole1*) following a modified protocol from Xiao^76^. Individual colonies were grown overnight at 30 °C in SD-media with glucose, lacking uracil. To cultivate Ole1 cells the media was supplemented with 0.02% linoleic acid (18:2 (n-6) and 0.2% Tween 60. To induce expression 0.5 ml from the overnight culture were transformed to 20 ml medium containing galactose and the appropriate supplements. Cells were cultured for 4 days at 30 °C, harvested by centrifugation at 3000 xg for 10 min, freezed at −20 °C and lyophilized for 48 hs.

***Picocyanobacterial cultivation.*** *Prochlorococcus* MED4 was grown in a seawater-based medium Pro99 medium^77^ based on Mediterranean seawater. *Synechococcus* WH8109 and WH7805 were grown in an artificial seawater-based medium^78^ with modifications as previously described^79^. All three strains were grown at 22 °C under cool white light under a 14:10h light-dark cycle, at a 30 μmol photonm^-2^s^-1^. The bacteria were harvested at the beginning of the stationary phase by centrifugation at 6000 ×g, 15 min, and then again at 10000 ×g for 10 min. Pellets were flash-freezed and stored in −80 °C until they were lyophilized for 24 hs.

***Lipid extraction and analysis.*** For analysis of esterified fatty acids in yeast, lyophilized cell pellets were submitted to transesterification using sodium methoxide^80^: Cells were homogenized in 0.5 ml 0.5 M sodium methoxide and 1.4 ml methanol by vortexing. After shaking for 1 h fatty acid methyl esters (FAME) were extracted by adding 2 ml saturated sodium chloride and 4 ml hexane. The hexane phase was dried under streaming nitrogen and dissolved in 30 μl acetonitrile.

For analysis of fatty acid profiles from cyanobacteria, lyophilized bacteria cells were submitted to acidic hydrolysis^81^. 1 ml of a methanolic solution containing 2.75% (v/v) sulfuric acid (95-97%) and 2% (v/v) dimethoxypropan was added to the sample. The sample was incubated for 1 h at 80 °C. To extract the resulting FAME, 200 μl of saturated sodium chloride solution and 2 ml of hexane were added. The hexane phase was dried under streaming nitrogen and dissolved in 100 μl acetonitrile for GC analysis.

For determination of the position of double bonds in fatty acids, fatty acid methyl esters (FAME) were converted into their 4,4-dimethyloxazoline (DMOX) derivatives according to Christie^82^. 90 μl of FAME resulting from acidic hydrolysis were dried under streaming nitrogen, 200 μl 2-amino-2-methyl-1-propanol was added and the sample was incubated at 180 °C for at least 14 h. Fatty acid derivatives were extracted by adding 1 ml of dichloromethane to the sample followed by 2.5 ml hexane and 1 ml water. The hexane phase was washed once with 1 ml water, then dried under streaming nitrogen. DMOX derivatives were separated from remaining FAME by TLC, using petrol ether/diethyl ether (2:1, v/v) as running solvent. DMOX derivatives were extracted from the plate, dissolved in 10 μl acetonitrile and subjected to GC/MS.

GC/FID analysis was performed with an Agilent (Waldbronn, Germany) 6890 gas chromatograph fitted with a capillary DB-23 column (30 m × 0.25 mm; 0.25 μm coating thickness; J&W Scientific, Agilent). Helium was used as carrier gas at a flow rate of 1 ml/min. The temperature gradient was 150 °C for 1 min, 150 – 200 °C at 8 K/min, 200–250 °C at 25 K/min and 250 °C for 6 min. GC/MS analysis for DMOX derivatives was carried out using a ThermoFinnigan Polaris Q mass selective detector connected to ThermoFinnigan Trace gas chromatograph equipped with a capillary DB-23 column. GC was performed using the same conditions as for GC/FID. Electron energy of 70 eV, an ion source temperature of 230 °C, and a temperature of 260 °C for the transfer line is used.

## Acknowledgements

We thank Debbie Lindell for kindly providing cyanobacterial strains, Andrea Nickel and Sabine Freitag for expert technical assistance and Cornelia Herrfurth for her advice with fatty acid analysis. This work was supported by funding from the People Programme (Marie Curie Actions) of the European Union’s Seventh Framework Programme FP7/2007-2013/ under REA grant agreement no. 317184, a European Commission ERC Advanced Grant no. 321647, and the Louis and Lyra Richmond Memorial Chair in Life Sciences (O.B.).

### Author Contributions

S.R. and O.B designed the project. S.R. conducted the molecular biology experiments, J.F.-U., and I.S. performed bioinformatics and E.H. and I.F. performed lipidomics; S.R. and O.B. wrote the manuscript with contributions from all authors to data analysis, figure generation, and the final manuscript.

### Competing interests

The authors declare no competing interests.

